# Morning glory species co-occurrence is associated with asymmetrically decreased and cascading reproductive isolation

**DOI:** 10.1101/767970

**Authors:** Kate L Ostevik, Joanna L Rifkin, Hanhan Xia, Mark D Rausher

## Abstract

Hybridization between species can affect the strength of the reproductive barriers that separate those species. Two extensions of this effect are: (1) the expectation that asymmetric hybridization will have asymmetric effects on reproductive barrier strength and (2) the expectation that local hybridization will affect only local reproductive barrier strength and could therefore alter within-species compatibility. We tested these hypotheses in a pair of morning glory species that exhibit asymmetric gene flow from highly selfing *Ipomoea lacunosa* into mixed mating *I. cordatotriloba* in regions where they co-occur. Because of the direction of this gene flow, we predicted that reproductive barrier strength would be more strongly affected in *I. cordatotriloba* than *I. lacunosa*. We also predicted that changes to reproductive barriers in sympatric *I. cordatotriloba* populations would affect compatibility with allopatric populations of that species. We tested these predictions by measuring the strength of a reproductive barrier to seed set across the species’ ranges. Consistent with our first prediction, we found that sympatric and allopatric *I. lacunosa* produce the same number of seeds in crosses with *I. cordatotriloba*, whereas crosses between sympatric *I. cordatotriloba* and *I. lacunosa* are more successful than crosses between allopatric *I. cordatotriloba and I. lacunosa.* This difference in compatibility appears to reflect an asymmetric decrease in the strength of the barrier to seed set in sympatric *I. cordatotriloba*, which could be caused by *I. lacunosa* alleles that have introgressed into *I. cordatotriloba*. We further demonstrated that changes to sympatric *I. cordatotriloba* have decreased its ability to produce seeds with allopatric populations of the same species, in line with our second prediction. Thus, in a manner analogous to cascade reinforcement, we suggest that introgression associated with hybridization not only influences between-species isolation but can also contribute to isolation within a species.

**Impact Statement:** Biological diversity depends on traits that prevent different species from successfully interbreeding. However, these reproductive barriers are often imperfect, leading to hybrid matings and possible genetic exchange between species where they occur together. When this happens, the reproductive barriers that separate species can themselves evolve to become stronger or weaker. Understanding the effects of hybridization on reproductive barriers is key to predicting the potential for future hybridization between species and ultimately whether hybridizing species will diverge, persist, or merge in regions where they co-occur. Here we hypothesize and show that hybridization in only one direction causes unidirectional changes to reproductive barrier strength and that geographically restricted hybridization causes local changes to barrier strength that can affect interbreeding within a species. Specifically, we found that gene flow from one species of morning glory into another likely caused a reproductive barrier to decrease in regions where they co-occur. The decreased reproductive barrier is caused by changes in only the species that received gene flow. We also found that the locally reduced barriers in the species that received gene flow affected reproductive compatibility between populations within that species. Thus, a breakdown of barriers between species can cause a build-up of barriers within a species. Our work demonstrates critical and rarely explored interactions at species boundaries.

## Introduction

Reproductive barriers are fundamental to the evolution and maintenance of biological diversity. They maintain the integrity of species by preventing hybridization and the homogenizing effect of gene flow. However, reproductive barriers are frequently incomplete, and closely related species often hybridize where they co-occur (Mallet 2005, Whitney et al. 2010). When hybridization occurs, such as in cases of secondary contact, it often leads to genetic exchange between species, and it can cause the strength of reproductive barriers to increase or decrease. These changes feed back to affect the potential for subsequent hybridization between the species. Whether reproductive isolation increases, decreases, or remains unchanged during hybridization determines if a pair of species in sympatry will collapse into a single species, complete speciation, or coexist in a stable hybrid zone (Endler 1977, Abbott et al. 2013, Todesco et al. 2016).

Hybridization increases or decreases reproductive isolation in two major ways: (1) by homogenizing genotypes and traits through gene flow and (2) by creating opportunities for selection. First, if hybridization homogenizes genotypes, and thus traits, that underlie reproductive barriers, the strength of reproductive isolation between species can increase or decrease. For example, when a trait that causes assortative mating on its own (e.g., self-fertilization) spreads to a new species, reproductive isolation will increase (Felsenstein 1981, Ortíz-Barrientos and Noor 2005). By contrast, homogenization of a trait that produces reproductive isolation through differentiation (e.g., flowering time) will cause isolation to decrease. Second, selection can act to change barriers in ways that depend on the fitness of hybrids. When hybrids are fit, selection may disfavor alleles that limit potential mates or interspecific fertilizations and thus reduce prezygotic reproductive barrier strength. However, when hybrids are unfit, selection can either act to weed out those unfit hybrids, purging incompatible alleles and reducing postzygotic isolation (Barton and Bengtsson 1986, Gavrilets 1997, Lemmon and Kirkpatrick 2006), or act to directly favor earlier-acting reproductive barriers that limit resources wasted on unfit offspring in a process called reinforcement (Dobzhansky 1940, Blair 1955, Howard 1993, Servedio and Noor 2003).

Which of these trajectories occurs depends on many factors, including (1) the strength, timing, and genetic architecture of the initial reproductive barriers, (2) the strength and type of natural selection, and, (3) the genetic variation available to selection (Clarke 1966, Felsenstein 1981, Barton and Hewitt 1985, Sanderson 1989, Marshall et al. 2002, Lemmon and Kirkpatrick 2006, Bank et al. 2012, Gompert et al. 2012, Lindtke and Buerkle 2015, Harrison and Larson 2016, Costa et al. 2020). Unfortunately, because these factors are challenging to quantify, it is difficult to predict whether reproductive isolation will increase or decrease in any particular contact zone. However, in cases of asymmetric or geographically restricted hybridization, we can make predictions about how reproductive isolation will change. Specifically, we expect that asymmetric hybridization will have asymmetric effects on reproductive barriers and that any local change to reproductive barriers due to geographically restricted hybridization may cause reproductive barriers to arise within species.

Asymmetric hybridization and gene flow occur in many organisms (Tiffin et al. 2001, Lowry et al. 2008, Todesco et al. 2016, Abbott 2017). These asymmetries arise under many circumstances, including when one of the two hybridizing species (1) is more common in regions of sympatry (Burgess et al. 2005), (2) is more successful at backcrossing with hybrid offspring (Ippolito et al. 2004), or (3) has a greater genetic load (Bierne et al. 2002, Kim et al. 2018, Pickup et al. 2019). In systems with asymmetric hybridization and gene flow, we expect that changes to reproductive barrier strength will also be asymmetric. This has been documented in cases of reinforcement (Noor 1995, Jaenike et al. 2006, Yukilevich 2012), but we expect it to be true regardless of whether reproductive barriers increase or decrease. In the most extreme cases with unidirectional hybridization and no cost of wasted pollen, we expect that only the species receiving pollen and/or gene flow will experience homogenization, selection on introgressed alleles, or reinforcing selection, and will thus have the potential to evolve in ways that change the strength of isolation.

When species that hybridize do not have completely overlapping ranges, changes to barrier strength in sympatric populations can create reproductive isolation between sympatric and allopatric populations of the same species. This phenomenon has been seen in some cases of reinforcement, where new or strengthened reproductive barriers in sympatry cause incompatibility between sympatric and allopatric populations of the species experiencing reinforcement (e.g., Hoskin et al. 2005, Jaenike et al. 2006, Kozak et al. 2015). This is known as “cascade reinforcement” or “cascade speciation” (Ortíz -Barrientos et al. 2009). However, any local change in reproductive barriers is likely to have cascading effects on reproductive isolation within species. For example, if gene flow homogenizes traits causing reproductive isolation through differentiation (e.g. flowering time divergence) in only sympatric populations of a species, sympatric and allopatric populations will become phenotypically mismatched and thus isolated. Similarly, if incompatibility alleles introgress from one species into some populations of another, those alleles will cause incompatibilities between populations that experience introgression and those that do not. Therefore, we expect that both local increases and local decreases in barrier strength can cause barriers to arise within species.

These two hypotheses have seldom been explicitly tested. We test both using a pair of morning glory species that are ideally suited to address these questions. *Ipomoea cordatotriloba* and *I. lacunosa* are sister species that have partially overlapping ranges and exhibit strongly asymmetric introgression from *I. lacunosa* into *I. cordatotriloba* in the regions where they co-occur (Rifkin et al. 2019a; see below for details). Furthermore, these species are strongly but not completely reproductively isolated by a barrier to seed set in which interspecific crosses produce few or no seeds (Martin 1970, Abel and Austin 1981, Diaz et al. 1996, Duncan and Rausher 2013b). This crossing barrier could manifest anywhere from pollen-pistil interactions to early embryo development, and its strength could evolve as a result of homogenization or selection. Although vigorous, hybrids between *I. cordatotriloba* and *I. lacunosa* tend to produce less pollen than the parental species (Abel and Austin 1981; Rifkin 2017). The existence of less-fertile hybrids means that reinforcing selection could act to increase the crossing barrier as it acts before maternal provisioning is complete (Coyne 1974, Hopkins 2013). Alternatively, selection could act to lessen the hybrid incompatibility and crossing barrier.

Here we assess seed set after crosses among sympatric and allopatric populations of *I. lacunosa* and *I. cordatotriloba* to determine whether species co-occurrence affects the strength of the crossing barrier. Given highly asymmetric introgression from *I. lacunosa* into *I. cordatotriloba*, we expect that if the crossing barrier evolves, change will be greater in sympatric populations of *I. cordatotriloba* than in sympatric populations of *I. lacunosa*. In addition, we expect that any change to the crossing barrier in sympatric populations of *I. cordatotriloba* will cascade to cause a reproductive barrier between sympatric and allopatric populations of *I. cordatotriloba*.

## Methods

### Species information

*Ipomoea lacunosa* and *I. cordatotriloba* (Convolvulaceae) are sister species (Muñoz-Rodríguez et al. 2018) that likely diverged between 1 and 1.6 million years ago (Carruthers et al. 2020). The two species have overlapping ranges in the southeastern United States, likely as a result of recent secondary contact that began between 78 and 1000 years ago (unpublished data), but only *I. lacunosa* occurs north of North Carolina into Canada and only *I. cordatotriloba* occurs south and west into more of Mexico (Fig. 1A). Both species produce many bisexual, self-compatible flowers that open for a single day (Fig. 1B-C). However, populations of *I. cordatotriloba* range from nearly complete outcrossing to nearly complete selfing, while all populations of *I. lacunosa* are highly selfing (all selfing rates ≥ 0.89, Duncan and Rausher 2013a). Accordingly, *I. lacunosa* exhibits many traits that are considered part of the “selfing syndrome” (Ornduff 1969, Sicard and Lenhard 2011), including small pale flowers, little nectar, and a low pollen:ovule ratio (Fig. 1C, McDonald et al. 2011, Duncan and Rausher 2013a, Rifkin et al. 2019b).

**Figure 1.**
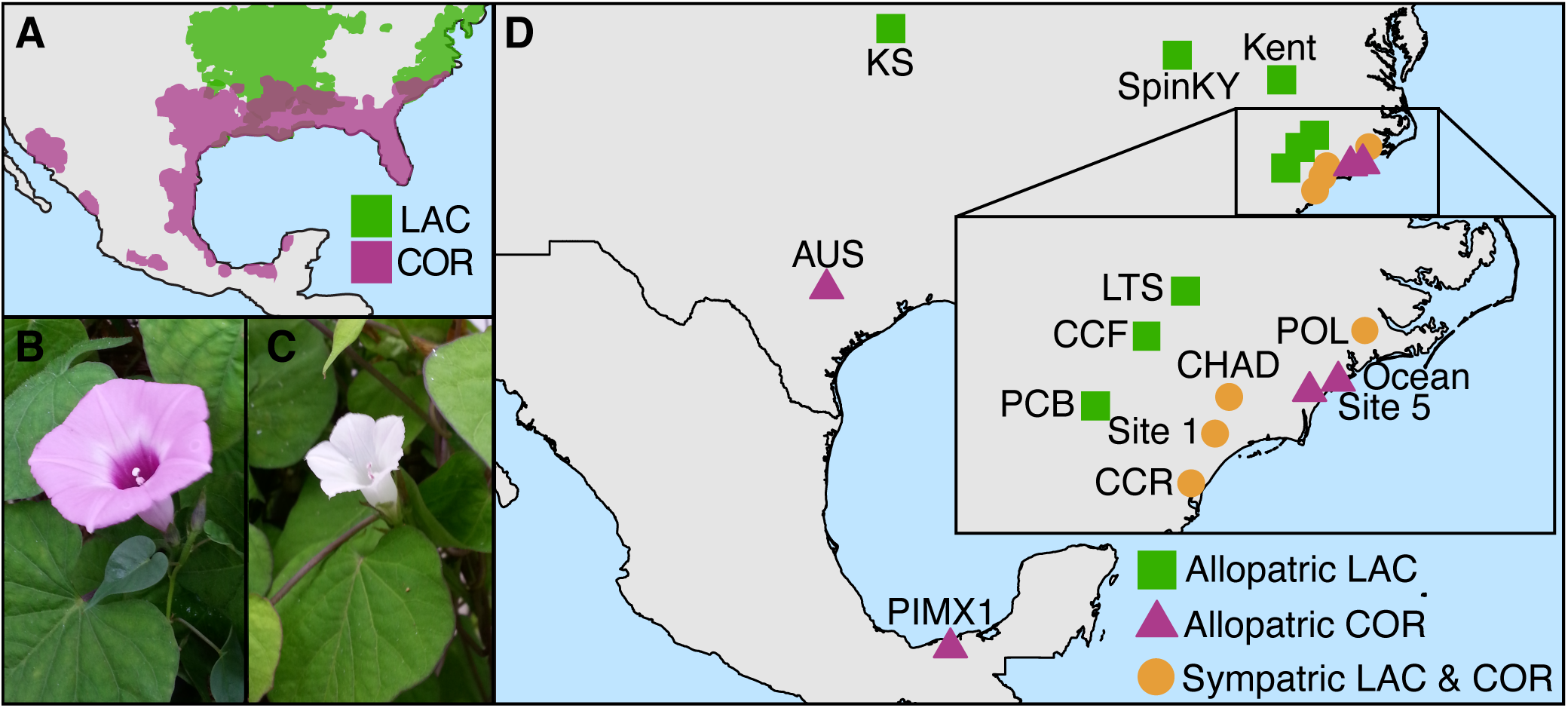
Information about our study species. A) Map showing the approximate distributions of *I. lacunosa* (LAC, green) and *I cordatotriloba* (COR, purple) in North America. This map was redrawn based on Khoury et al. (2015) and generally matches collection records from the United States (USDA, NRCS 2019). B) A typical allopatric *I. cordatotriloba* flower. C) A typical *I. lacunosa* flower. D) Map showing the locations of populations used in this study (see Table S1 for details).

Rifkin et al. (2019a) identified asymmetric gene flow from highly selfing *I. lacunosa* into mixed-mating *I. cordatotriloba*. Multiple genetic analyses revealed that *I. lacunosa* is genetically similar across its range, while *I. cordatotriloba* consists of two distinct genetic groups that correspond to allopatric and sympatric populations (Rifkin et al. 2019a). Genetic similarity between sympatric *I. cordatotriloba* and *I. lacunosa*, as measured either by π or allele-frequency differences, is substantially higher than between allopatric populations of the two species (Rifkin et al. 2019a). Moreover, sympatric populations of *I. cordatotriloba* contain alleles that are present in *I. lacunosa* but absent from allopatric *I. cordatotriloba*, often at frequencies greater than 0.5. However, the reverse is not common (Rifkin et al. 2019a). These results indicate that there has been substantial introgression from *I. lacunosa* into *I. cordatotriloba* in regions of sympatry in the recent past, while there has been essentially no introgression from *I. cordatotriloba* into *I. lacunosa.* Although introgression from a selfer into a mixed mater is counter to some expectations (Davis and Heywood 1963), a recent survey of gene flow between self-compatible species with different levels of selfing found more gene flow from selfing species into outcrossing species in every study examined (Pickup et al. 2019). One of several potential explanations for this pattern is that prior self-fertilization excludes fertilization by interspecific and hybrid pollen in selfing species (Lloyd 1979, Fishman and Wyatt 1999, Goodwillie and Weber 2018, Brys et al. 2016). In these *Ipomoea* species, anthers often dehisce before flowers open, so pre-emptive self-fertilization in *I. lacunosa* (where anthers and stigmas touch) is likely to explain at least some of the asymmetry in gene flow.

### Populations and crosses

To measure variation in the success of crosses made within and between *I. lacunosa* and *I. cordatotriloba* individuals, we grew 60 accessions from 18 populations collected across 14 locations (Fig. 1D, Table S1) under common greenhouse conditions (see Rifkin et al 2019b for details; these are a subset of the plants used for Qst estimation in that study). Briefly, seeds were scarified and germinated in soil in a growth room under a long-day cycle (16:8 light:dark) and shifted to a short-day cycle (12:12 light:dark) after approximately four weeks. When flower buds appeared, the plants were transferred to the Duke University Greenhouse Facility. We selected 40 focal plants made up of 10 individuals from each of the following four population categories: allopatric *I. cordatotriloba*, allopatric *I. lacunosa*, sympatric *I. cordatotriloba*, and sympatric *I. lacunosa*. Allopatric populations were defined as populations growing outside the range of the other species or populations within the range of the other species where we observed only one species after a thorough search. Data from Rifkin et al. (2019a) suggests that these categorizations capture biologically relevant groups.

We reciprocally crossed 6 flowers from each focal plant and a representative of each population category, so that each focal plant was involved in 48 crosses (2 directions x 4 categories x 6 crosses), 24 as a pollen recipient and 24 as a pollen donor (Fig. S1). Within each cross type (i.e., all pairwise combinations of the population categories), we made crosses between individuals from at least three populations within each population category, and we did not cross any combination of populations more than three times (Table S1). To perform each of the 1,920 crosses, we emasculated flowers the day before they opened by dissecting the corolla and removing the anthers with forceps. Between 8:00am and 12:00pm the next day, we pollinated the emasculated flowers by dabbing a flower from the individual used as the male on the stigma of the flower used as the female. This method transfers many more pollen grains than needed to fertilize the 4 ovules present in each pistil. In these species, if a flower is left unpollinated or a cross fails, the flower generally abscises and falls off the plant within 4 days. Therefore, we checked the crosses every day until they abscised or until the seed capsule had dried and the sepals had reflexed, indicating seed maturation. Finally, we counted and weighed the mature seeds and fruits.

### Statistical Analyses

To determine whether cross type affected whether a cross was successful, we used R version 3.4.2 (R Core Team 2020) and the R packages *lme4* (Bates et al. 2015) and *glmmTMB* (Brooks et al. 2017) to fit mixed effect models to two measures of cross success: (1) whether a fruit produced at least one mature seed (where mature seeds were defined as those with seed weight > 10 mg, which is consistent with our observations about seed viability) and (2) the mean number of mature seeds produced by a specific pair of individuals). All models included maternal individual nested within maternal population as a random effect. Models of fruit set included a binomial link and individual cross as a random effect, and models of mean seed number from interspecific crosses included a term for zero-inflation (see Table S2 for exact model specifications). We also fit models with paternal individual nested within paternal population as a random effect. However, paternal identity was not a significant model term and did not qualitatively change our results, and thus it was excluded from the results presented below. To determine whether geographic distance affected intraspecific cross success we fit a model to the mean number of seeds produced (as described above) that also included the distance between the populations of the individuals crosses as a factor. In all cases, we identified significant model terms using Wald chi-squared tests implemented in the R package *car* (Fox and Weisberg 2019) and significant contrasts using least-squared means implemented in the R package *emmeans* (Lenth 2016).

Although all the plants appeared to make healthy and functional flowers, we found that individuals from one allopatric *I. cordatotriloba* population, Ocean, were consistently less fertile than individuals from other populations (only 9% of intraspecific crosses set seed compared to 46-82% of intraspecific crosses in other populations). We removed this population from the analyses below, but our results do not qualitatively change when this population is included (Table S2).

## Results

Controlled crosses revealed variation in the success of different cross types. First, our results confirm the existence of a strong reproductive barrier separating *I. cordatotriloba* and *I. lacunosa*. Overall, 68% of intraspecific crosses and only 5% of interspecific crosses set fruit with at least one mature seed (Fig. 2). Interspecific crosses are significantly less successful than both crosses within *I. cordatotriloba* (Table S2; fruit set: z = 14.7, p < 0.001; mean seed number: t = 4.77, df = 295, p < 0.001) and crosses within *I. lacunosa* (Table S2; fruit set: z = 15.3, p < 0.001; mean seed number: t = 5.91, df = 295, p < 0.001). However, we found no evidence that the success of interspecific crosses is affected by which species is used as the maternal parent (Fig. 2; Table S2; fruit set: z = -1.12, p = 0.26; mean seed number: t = 0.84, df = 295, p = 0.40).

**Figure 2.**
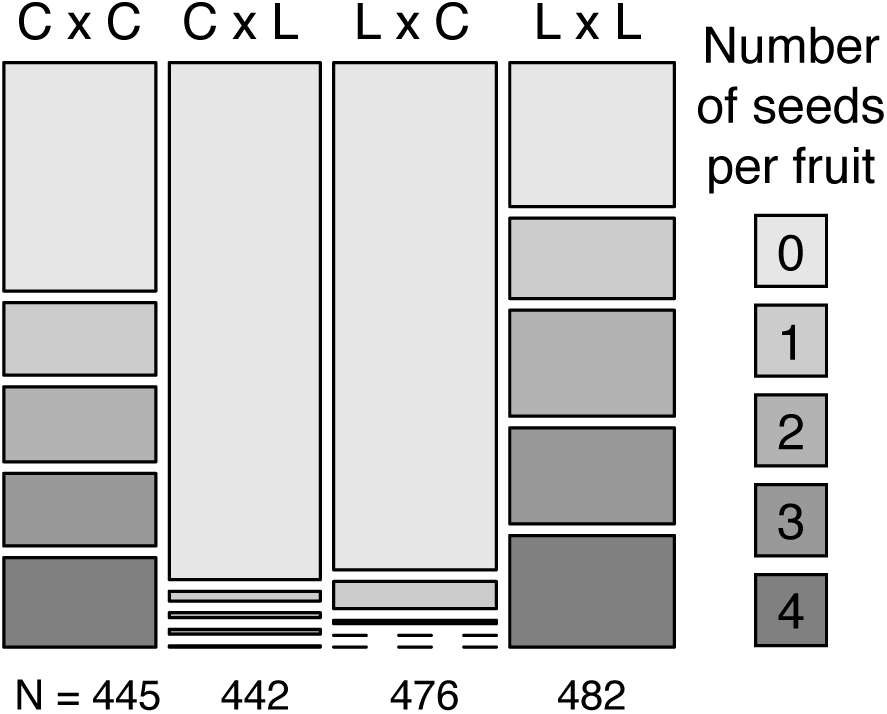
The effect of species on seed set. The number of seeds produced by different types of crosses (ovule parent x pollen parent; C = *I. cordatotriloba*, L = *I. lacunosa*). Each box is proportional to the number of fruits that contained indicated number of seeds after pollination. Dashed lines represent cases in which there were no fruits with a particular seed number.

Second, interspecific crosses made with plants from sympatric sites were significantly more likely to be successful than those made with plants from allopatric sites (Fig. 3: fruit set: z = -1.97, p = 0.048; mean seed number: t = -2.52, df = 146, p = 0.013). To determine whether differences in one or both species explain the pattern of higher seed set in sympatric plants, we compared the success rates of interspecific crosses using sympatric and allopatric plants within each species. We found that crosses between sympatric *I. cordatotriloba* and range-wide *I. lacunosa* are more successful than between allopatric *I. cordatotriloba* and range-wide *I. lacunosa* (Fig. 3; Table S2; fruit set: z = -1.97, p = 0.048; mean seed number: t = -3.81, df = 146, p < 0.01), whereas crosses between sympatric *I. lacunosa* and range-wide *I. cordatotriloba* are no more likely to be successful than those between allopatric *I. lacunosa* and range-wide *I. cordatotriloba* (Fig. 3; Table S2; fruit set: z = 0.10, p = 0.92; mean seed number: t = 0.28, df = 146, p = 0.78). These comparisons indicate that the higher seed set of interspecific crosses involving sympatric plants is due primarily to higher seed set in crosses involving sympatric *I. cordatotriloba.*

**Figure 3.**
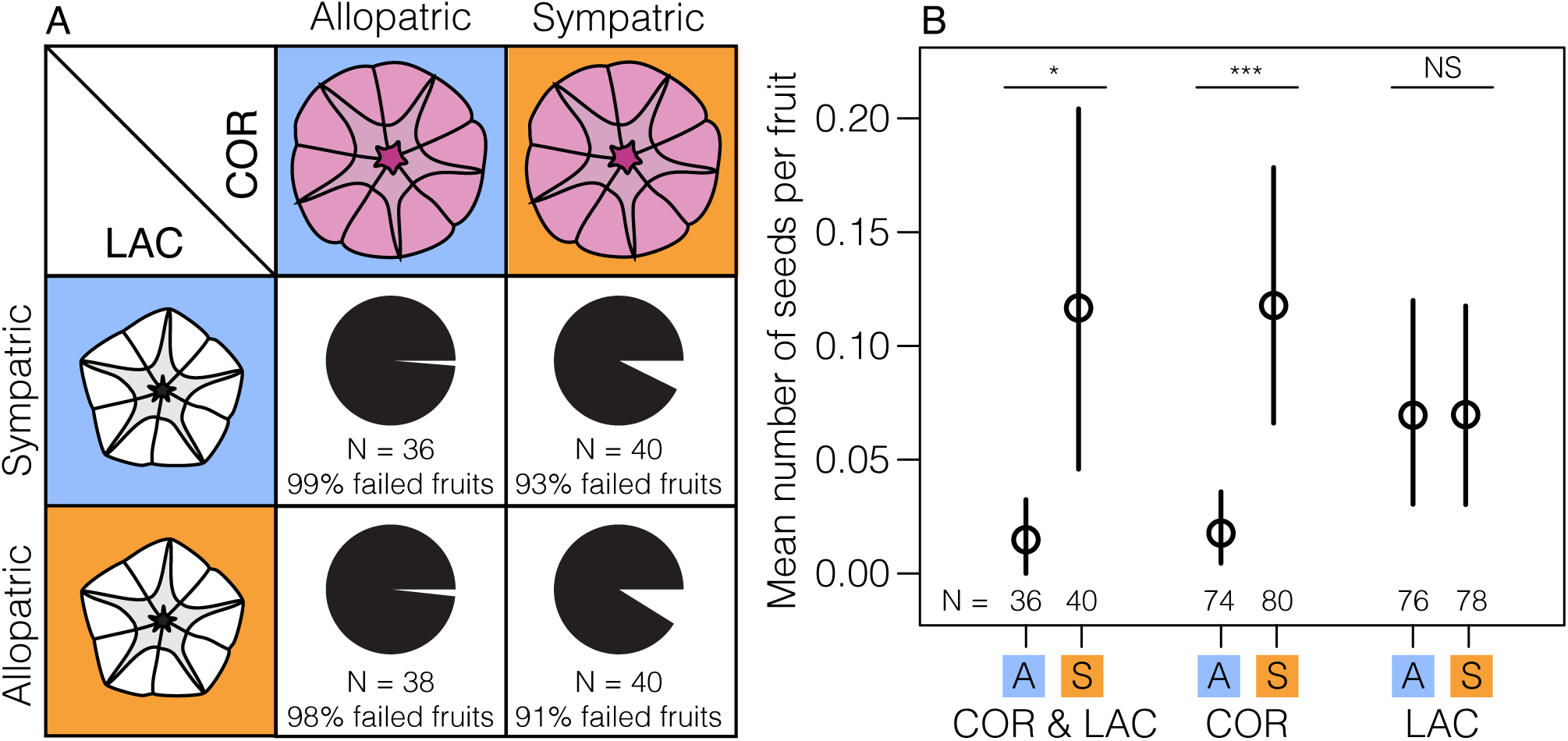
Interspecific cross success depends on species co-occurrence. A) The proportion of interspecific crosses that set at least one mature seed. B) The mean and 95% confidence intervals (based on bootstrap percentiles) for the mean number of seeds produced by different types of interspecific crosses (from left to right): both species are allopatric (A), both species are sympatric (S), allopatric *I. cordatotriloba* (COR) and any *I. lacunosa* (LAC), sympatric *I. cordatotriloba* and any *I. lacunosa*, allopatric *I. lacunosa* and any *I. cordatotriloba*, sympatric *I. lacunosa* and any *I. cordatotriloba*.

Finally, we found that crosses made within *I. cordatotriloba* were affected by whether the individuals were from allopatric or sympatric populations (Fig. 4A; Table S2; fruit set: = 14, df = 2, p = 0.0011; mean seed number: = 8.5, df = 2, p = 0.014), while this was not true for crosses within *I. lacunosa* (Fig. 4B; Table S2; fruit set: = 1.0, df = 2, p = 0.60; mean seed number: : = 0.41, df = 2, p = 0.82). Within *I. cordatotriloba*, crosses between two allopatric populations were more successful than crosses between allopatric and sympatric populations (Fig. 4; Table S2; fruit set: z = 3.7, p < 0.001; mean seed number: t = 2.8, df = 54, p = 0.019) and were marginally more successful than crosses between two sympatric populations (Fig. 4; Table S2; fruit set: z = 2.8, p = 0.013; mean seed number: t = 2.1, df = 54, p = 0.094). This variation within *I. cordatotriloba* does not appear to be caused by population variation correlated with geographic distance because geographic distance is not significantly correlated with intraspecific cross success in either *I. cordatotriloba* (Fig S3; fruit set: = 1.9, df = 1, p = 0.17; mean seed number: : = 2.61, df = 1, p = 0.10) or *I. lacunosa* (Fig S3; fruit set: = 0.28, df = 1, p = 0.60; mean seed number: = 0.038, df = 1, p = 0.85). In fact, there is a weak trend in the direction opposite to this expectation, where *I. cordatotriloba* individuals from more distance populations tend to be more compatible (Fig S3).

**Figure 4.**
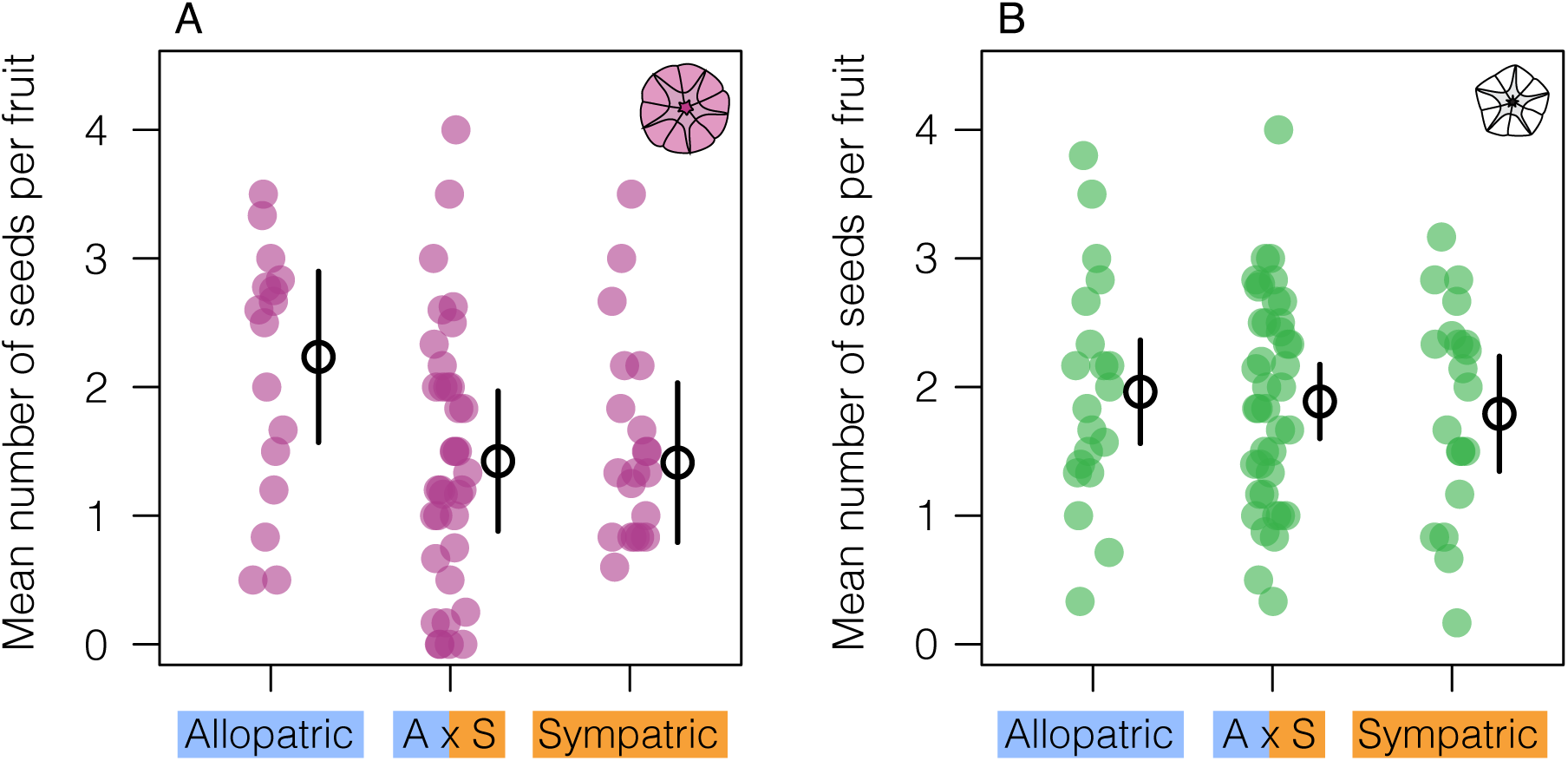
The success of crosses within I. *cordatotriloba*, but not within *I. lacunosa*, depends on species co-occurrence. The mean and 95% confidence intervals for the mean number of seeds produced by crosses within *I. cordatotriloba* (A) and within *I. lacunosa* (B) where the individuals crossed are from different population types (from left to right): both are from allopatric populations, one is from an allopatric population and the other is from a sympatric population, both are from sympatric populations. Each point represents the mean number of seeds produced by 5-6 crosses made with same individuals.

## Discussion

We have shown that *I. cordatotriloba* and *I. lacunosa* are separated by a barrier to seed set that is weaker in regions where the two species occur together. This result could explain the wide variation in crossing success reported in previous studies (Martin 1970, Abel and Austin 1981, Diaz et al. 1996, Duncan and Rausher 2013b). The weaker barrier appears to be caused by changes in the sympatric populations of *I. cordatotriloba* and not *I. lacunosa*, a result consistent with our hypothesis that asymmetric hybridization and gene flow will lead to asymmetric change in barrier strength. Further, the weaker barrier in sympatric *I. cordatotriloba* likely explains a crossing barrier that we observe between sympatric and allopatric individuals of *I. cordatotriloba.* This result is consistent with our second hypothesis that any local change to reproductive barriers can cause barriers to arise within a species.

Variable reproductive barrier strength has often been associated with variable environmental conditions (e.g., Mecham 1960, Grant and Grant 1993, Seehausen et al 1997, Taylor et al. 2006, Mercander et al. 2009, Gilman et al. 2011). However, because we measured the strength of cross-compatibility in a common garden, environmental variation is unlikely to explain the variation in barrier strength we observe here. Instead, the association between species co-occurrence and a weaker reproductive barrier suggests that hybridization and gene flow have eroded the strength of the crossing barrier. This is noteworthy because, while the erosion of barriers in sympatry is supported by theory (Barton and Hewitt 1985, Butlin 1987, Virdee and Hewitt 1994, Kelly and Noor 1996) and is implied by patterns of genetic variation in many systems (e.g., Bettles et al. 2005, Borge et al 2005, Gow et al. 2006, Sander et al. 2014, Zuellig and Sweigart 2019), studies in which the strength of reproductive barriers have been explicitly shown to be weaker in sympatry are rare (but see Virdee and Hewitt 1994, Saetre et al. 1999, Carney et al. 2000, Collins and Rawlins 2013). Although it would be interesting to know why the strength of the crossing barrier decreased rather than increased, this study does not allow us to address this issue. However, it is possible that the asymmetric hybridization and gene flow in this system partially explains why the crossing barrier has not been reinforced as Servedio and Kirkpatrick (1997) showed theoretically that reinforcement is less likely in cases with one-way migration relative to symmetric migration.

The weaker crossing barrier could be explained by the homogenization of *I. cordatotriloba* and *I. lacunosa* in sympatry. Hybrids produced by controlled crosses between the species are more cross-compatible with the parental types than the parents are with each other (unpublished data) suggesting that genetic similarity is associated with cross-compatibility in this system. Further, the asymmetric erosion of the crossing barrier is consistent with asymmetric introgression of alleles that underlie the barrier from *I. lacunosa* into *I. cordatotriloba*. For example, consider a scenario in which the crossing barrier is caused by incompatible alleles in zygotes (such that zygotes are unlikely to develop into seeds) at a locus that has allele *A* in allopatric *I. cordatotriloba* populations and allele *a* in allopatric *I. lacunosa* populations. When both gametes in a cross carry the same allele, there is no incompatibility, while if one gamete is *A* and the other is *a*, there is incompatibility. When secondary contact first occurs, within-species matings are compatible and between-species matings are incompatible. After asymmetric introgression, however, the frequency of *a* in the sympatric *I. cordatotriloba* population is high. Therefore, a large proportion of the crosses between this population and sympatric *I. lacunosa* will involve both gametes carrying allele *a*, which are compatible. The crossing barrier between sympatric *I. cordatotriloba* and *I. lacunosa* will thus be lower than between allopatric populations of the two species. We note that this argument is easily extended to loci that underlie an earlier failure of the cross (e.g., loci that cause a mismatch between pollen and pistils or pollen and ovules) or to multiple loci, each of which contributes incrementally to incompatibility.

Interestingly, our finding that a crossing barrier somewhat isolates allopatric and sympatric populations within *I. cordatotriloba*, but not within *I. lacunosa*, is also consistent with the asymmetric introgression of alleles that underlie the barrier from *I. lacunosa* into *I. cordatotriloba*. After introgression in the scenario described above, sympatric *I. cordatotriloba* populations would harbor *a* alleles from *I. lacunosa* and would be incompatible with allopatric populations of *I. cordatotriloba* that have A alleles. At the same time, sympatric I. *lacunosa* populations would still be fixed for *a* alleles and would not be incompatible with their allopatric counterparts. Furthermore, in the case where the introgression of *a* alleles into sympatric *I. cordatotriloba* populations was not complete, A and a alleles would be segregating in the sympatric populations and could cause crosses made between two sympatric *I. cordatotriloba* individuals to also fail. This could explain the marginally reduced number of seeds set after crosses made between sympatric *I. cordatotriloba* in this study.

Although the above explanation is compelling, it is possible that processes other than introgression explain the pattern of asymmetric change. First, it could be that *I. cordatotriloba* simply has greater population variation in reproductive barrier strength that happens to correlate with sympatric regions. However, this seems unlikely given that geographic variation does not explain cross success. Second, it could be that a lack of genetic diversity present in *I. lacunosa* (Rifkin 2019a) prevented the evolution of reproductive barriers in that species. In a scenario where there is direct selection on reproductive isolation in sympatry, it is possible that only *I. cordatotriloba* harboured the genetic variation needed to respond. A lack of genetic variation for stronger reproductive barriers is often cited in studies that fail to find reinforcement (Barton and Hewitt 1985). However, it is also theoretically possible in cases where selection favors weaker barriers. Future studies that determine the precise nature and genetic basis of the crossing barrier will allow us to continue to evaluate the merit of these alternative explanations.

It is possible that the asymmetric erosion of the crossing barrier seen here could cause a positive feedback in which that erosion facilitates even more asymmetric gene flow that eventually leads to the extinction of *I. cordatotriloba*. However, it is important to remember that we are only tracking the effect of hybridization on a single reproductive barrier. Other reproductive barriers (e.g., flowering time and pollinator isolation) could be increasing or decreasing. Indeed, it seems that reproductive isolation caused by differences in the rate of self-fertilization may have increased in sympatry. Rifkin et al. (2019a) observed phenotypic convergence between sympatric *I. cordatotriloba* and *I. lacunosa* caused by change in *I. cordatotriloba* but little change in *I. lacunosa* for anther-stigma separation (herkogamy) and selfing rate. Because herkogamy is the main determinant of selfing rate in self-compatible *Ipomoea* (Duncan and Rausher 2013a) and other plant species (e.g., Karron et al. 1997, Motten et al. 2000), the significantly reduced anther-stigma separation in sympatric *I. cordatotriloba* plants is consistent with the substantially higher selfing rates observed in this species in sympatric regions. This increased selfing in *I. cordatotriloba* presumably constitutes a barrier to gene flow from *I. lacunosa*, but this needs to be confirmed by additional experiments. If true, however, the decreased reproductive isolation caused by decreased cross incompatibility we document here may be offset, at least to some extent, by an increase in isolation caused by the effects of gene flow on selfing rate. Interestingly, this may also partly explain why the crossing barrier was eroded and not reinforced. Simulations by Castillo et al. (2016) showed that increases in self-compatibility in sympatry often preclude the evolution of other assortative mating traits (e.g., a stronger crossing barrier). Either way, our study suggests that not all reproductive barriers will respond to hybridization in the same way.

Together, the intraspecific crossing barrier and increased selfing in regions of sympatry suggest that the allopatric and sympatric populations of *I. cordatotriloba* are substantially reproductively isolated, consistent with the genetic differentiation observed in (Rifkin et al 2019a). Because both of these effects appear to be by-products of introgression in sympatry, our results provide an example of how asymmetric introgression in sympatry has the potential to drive speciation between allopatric and sympatric populations of the same species. This process is analogous to cascade reinforcement (Ortíz -Barrientos et al. 2009) except that erosion, rather than reinforcement, of the between-species crossing barrier causes an increase in isolation between sympatric and allopatric populations. Our work thus demonstrates that any type of evolutionary change in local reproductive barriers has the potential to affect within-species compatibility.

## Acknowledgements

We would like to thank Evangeline Mareki, Irene Liao, and Yuncheng Duan for help making crosses and managing morning glories. We would also like to thank Jenn Coughlan, Kieran Samuk, Felix Beaudry, Loren Rieseberg, Spencer Barrett, Stephen Wright, Elizabeth Lacey, the Rausher lab, and three anonymous reviewers for helpful discussions and comments on earlier versions of this manuscript. This research was supported by a National Science and Engineering Research Council of Canada Postdoctoral Fellowship (516658) to KLO, a National Science Foundation Doctoral Dissertation Improvement Grant (DEB 1501954) to JLR, and a National Science Foundation grant (DEB 1542387) to MDR.

## Author Contributions

KLO, JLR, and MDR designed the study. KLO and HX collected the data. KLO analyzed the data. KLO, JLR, and MDR drafted the manuscript. All authors read, edited, and approved the final manuscript.

## Data Accessibility

The data and scripts used in these analyses will be made available on an online data repository (Dryad) upon manuscript acceptance.

## Notes

### Competing Interest Statement

The authors have declared no competing interest.

