## Supplemental Material for "Morning glory species co-occurrence is associated with asymmetrically decreased and cascading reproductive isolation"

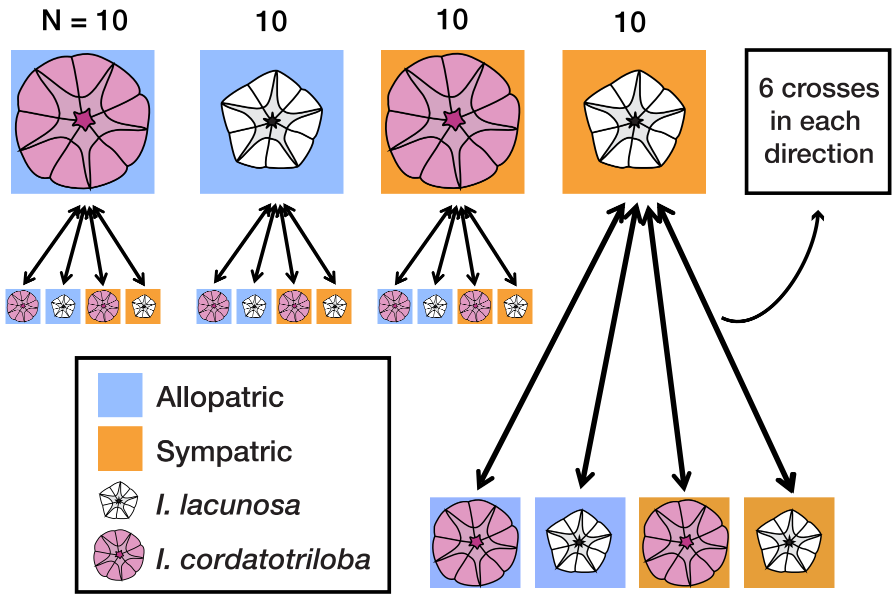


Figure S1 – Diagram of the full crossing design for this study.

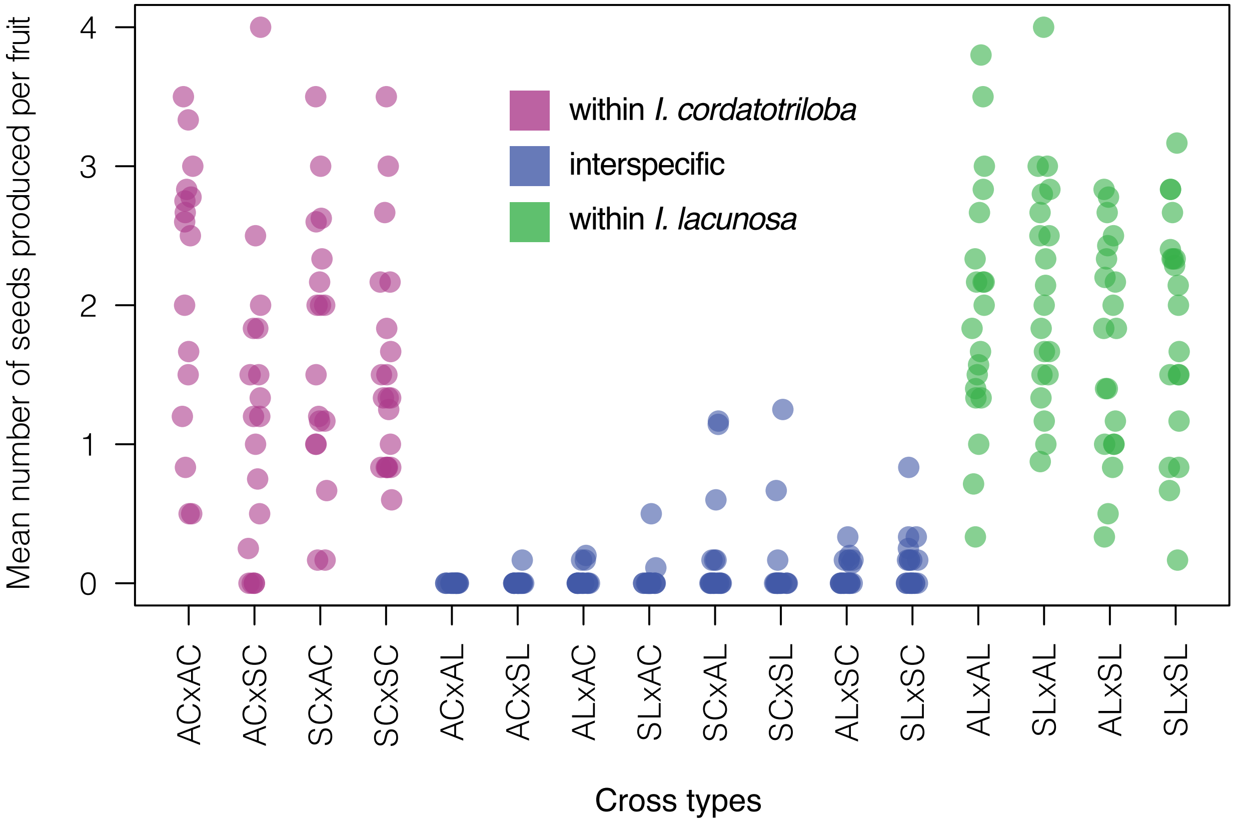


Figure S2 – Cross type affects mean seed number. The mean number of seeds produced by crosses between all pairwise combinations of population categories: allopatric *I. cordatotriloba* (AC), allopatric *I. lacunosa* (AL), sympatric *I. cordatotriloba* (SC), and sympatric *I. lacunosa* (SL). Each point represents 5-6 crosses.

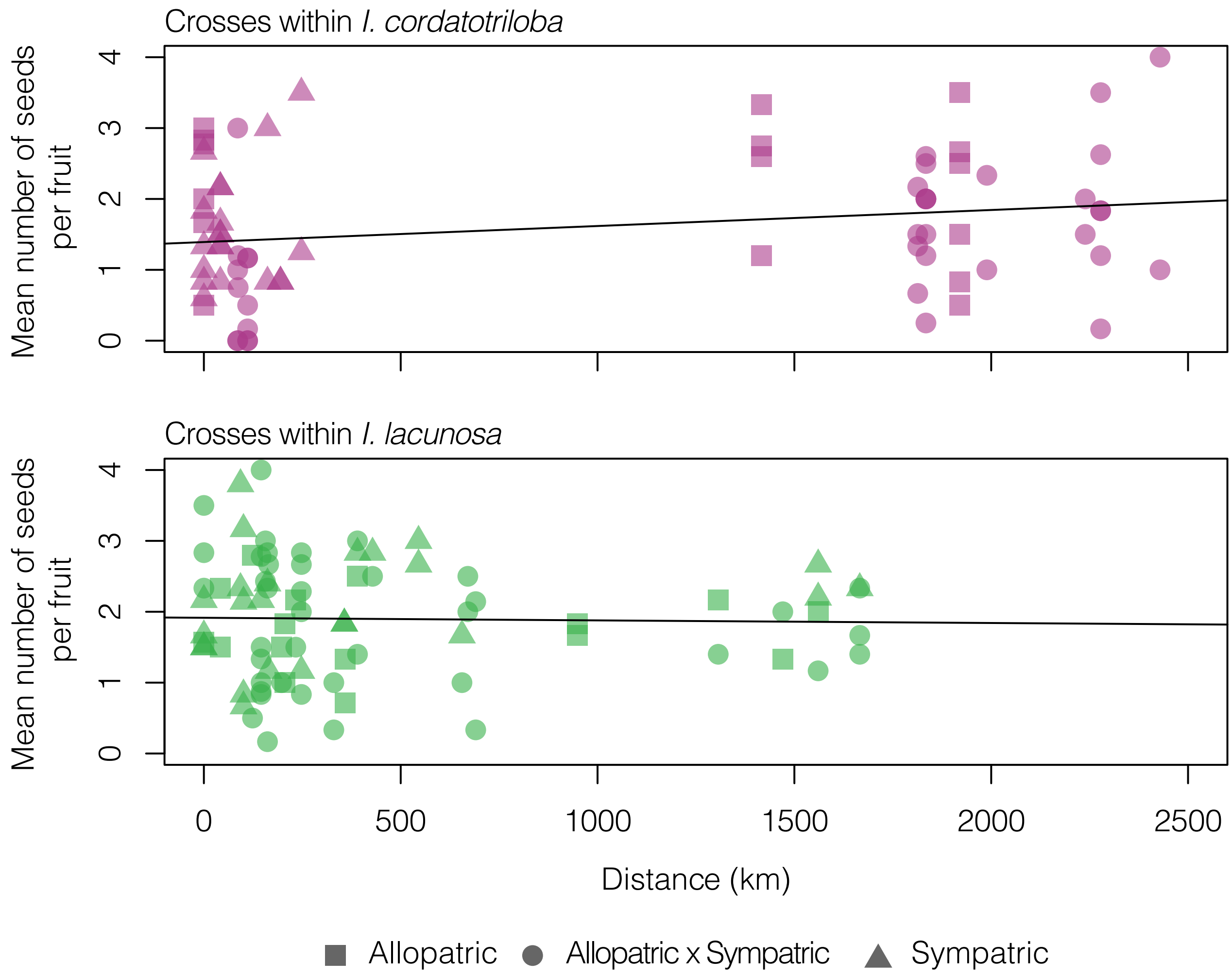


Figure S3 – Cross success is not affected by geographic distance. The mean number of seeds produced by crosses within *I. cordatotriloba* and *I. lacunosa* versus the geographic distance between where the individuals were collected. Each point represents 5-6 crosses, and shape represented whether the individuals crossed were both from allopatric populations, both from sympatric populations, or from one of each population type. Distance does not significantly affect cross success in *I. cordatotriloba* (: = 2.61, df = 1, p = 0.10) or *I. lacunosa* (: = 0.038, df = 1, p = 0.85).

Table S1 – Information about the accessions and populations used in this study. Source codes: RL = Rausher Lab, USDA = United States Department of Agriculture, BL = Baskin Labs, OP = seed collected from cuttings brought into the lab, S = selfed seed collected from plants grown in the greenhouse from seed.

| ID | Species | Site | Accession | Source | Latitude | Longitude | Geography |
|---|---|---|---|---|---|---|---|
| 11 | COR | AUS | AUS_PC_2 | RL-OP | 30.25550 | -97.70958 | Allopatric |
| 14 | COR | AUS | AUS_PC_8 | RL-OP | 30.25550 | -97.70958 | Allopatric |
| 38 | COR | AUS | AUS_PC_4 | RL-OP | 30.25550 | -97.70958 | Allopatric |
| 44 | COR | AUS | AUS_PC_8 | RL-OP | 30.25550 | -97.70958 | Allopatric |
| 55 | COR | AUS | AUS_PC_2 | RL-OP | 30.25550 | -97.70958 | Allopatric |
| 12 | LAC | CCF | CCF_L_2-1 | RL-S | 34.89755 | -79.8021 | Allopatric |
| 16 | LAC | CCR | CCR_L_1-2 | RL-S | 33.48225 | -79.27 | Sympatric |
| 22 | LAC | CCR | CCR_L_2-2 | RL-S | 33.48225 | -79.27 | Sympatric |
| 32 | LAC | CCR | CCR_L_1-2 | RL-S | 33.48225 | -79.27 | Sympatric |
| 34 | COR | CCR | CCR_PC_2-1 | RL-S | 33.48225 | -79.27 | Sympatric |
| 100 | LAC | CCR | CCR_L_2-2 | RL-S | 33.48225 | -79.27 | Sympatric |
| 103 | LAC | CCR | CCR_L_3-1 | RL-S | 33.48225 | -79.27 | Sympatric |
| 3 | LAC | CHAD | CHAD_L_3-c1 | RL-S | 34.31021 | -78.8264 | Sympatric |
| 10 | LAC | CHAD | CHAD_L_2-1 | RL-S | 34.31021 | -78.8264 | Sympatric |
| 40 | COR | CHAD | CHAD_PC_3-2 | RL-S | 34.31021 | -78.8264 | Sympatric |
| 68 | COR | CHAD | CHAD_PC_2-2 | RL-S | 34.31021 | -78.8264 | Sympatric |
| 83 | COR | CHAD | CHAD_PC_1-1 | RL-S | 34.31021 | -78.8264 | Sympatric |
| 87 | COR | CHAD | CHAD_PC_1-1 | RL-S | 34.31021 | -78.8264 | Sympatric |
| 93 | LAC | CHAD | CHAD_L_3-c1 | RL-S | 34.31021 | -78.8264 | Sympatric |
| 102 | COR | CHAD | CHAD_PC_3-2 | RL-S | 34.31021 | -78.8264 | Sympatric |
| 105 | COR | CHAD | CHAD_PC_2-2 | RL-S | 34.31021 | -78.8264 | Sympatric |
| 110 | LAC | CHAD | CHAD_L_4-1 | RL-S | 34.31021 | -78.8264 | Sympatric |
| 28 | LAC | Kent | Kent_L_8-1 | RL-S | 37.19345 | -80.5737 | Allopatric |
| 41 | LAC | Kent | Kent_L_7-1 | RL-S | 37.19345 | -80.5737 | Allopatric |
| 46 | LAC | Kent | Kent_L_8-1 | RL-S | 37.19345 | -80.5737 | Allopatric |

| | | | | | | | |
|---|---|---|---|---|---|---|---|
| 58 | LAC | Kent | Kent_L_6-1 | RL-S | 37.19345 | -80.5737 | Allopatric |
| 29 | LAC | KS | KS_L_5-1 | RL-S | 38.88445 | -95.3206 | Allopatric |
| 37 | LAC | KS | KS_L_4-2 | RL-S | 38.88445 | -95.3206 | Allopatric |
| 97 | LAC | KS | KS_L_8-1 | RL-S | 38.88445 | -95.3206 | Allopatric |
| 5 | LAC | LTS | LTS_L_6-c1 | RL-S | 35.33599 | -79.3535 | Allopatric |
| 80 | LAC | LTS | LTS_L_5-1 | RL-S | 35.33599 | -79.3535 | Allopatric |
| 90 | LAC | LTS | LTS_L_6-c1 | RL-S | 35.33599 | -79.3535 | Allopatric |
| 150 | COR | Ocean | Ocean 4 | RL-OP | 34.48462 | -77.5637 | Allopatric |
| 9 | LAC | PCB | PCB_L_1-2 | RL-S | 34.22187 | -80.4021 | Allopatric |
| 21 | LAC | PCB | PCB_L_6-2 | RL-S | 34.22187 | -80.4021 | Allopatric |
| 23 | LAC | PCB | PCB_L_2-1 | RL-S | 34.22187 | -80.4021 | Allopatric |
| 63 | LAC | PCB | PCB_L_6-2 | RL-S | 34.22187 | -80.4021 | Allopatric |
| 30 | COR | PIMX1 | PI 518494 01 SD_PC_MX_71-1 | USDA-S | 18.2 | -93.0831 | Allopatric |
| 62 | COR | PIMX1 | PI 518494 01 SD_PC_MX_71-1 | USDA-S | 18.2 | -93.0831 | Allopatric |
| 64 | COR | PIMX1 | PI 518494 01 SD_PC_MX_44-c2 | USDA-S | 18.2 | -93.0831 | Allopatric |
| 73 | COR | PIMX1 | PI 518494 01 SD_PC_MX_44-c2 | USDA-S | 18.2 | -93.0831 | Allopatric |
| 13 | COR | POL | POL_PC_6-2 | RL-S | 34.94662 | -77.241 | Sympatric |
| 19 | COR | POL | POL_PC_6-2 | RL-S | 34.94662 | -77.241 | Sympatric |
| 57 | LAC | POL | POL_L_4-1 | RL-S | 34.94662 | -77.241 | Sympatric |
| 94 | LAC | POL | POL_L_3-2 | RL-S | 34.94662 | -77.241 | Sympatric |
| 101 | LAC | POL | POL_L_4-1 | RL-S | 34.94662 | -77.241 | Sympatric |
| 117 | LAC | POL | POL_L_3-2 | RL-S | 34.94662 | -77.241 | Sympatric |
| 27 | COR | Site1 | Site 1_PC_3-1 | RL-S | 33.95844 | -78.9915 | Sympatric |
| 42 | LAC | Site1 | Site 1_L_1-1 | RL-S | 33.95844 | -78.9915 | Sympatric |
| 43 | LAC | Site1 | Site 1_L_1-1 | RL-S | 33.95844 | -78.9915 | Sympatric |
| 99 | COR | Site1 | Site 1_PC_3-1 | RL-S | 33.95844 | -78.9915 | Sympatric |
| 104 | COR | Site1 | Site 1_PC_2-1 | RL-S | 33.95844 | -78.9915 | Sympatric |
| 108 | COR | Site1 | Site 1_PC_2-1 | RL-S | 33.95844 | -78.9915 | Sympatric |
| 24 | COR | Site5 | Site 5_PC_1-1 | RL-S | 34.37725 | -77.8966 | Allopatric |
| 75 | COR | Site5 | Site 5_PC_1-1 | RL-S | 34.37725 | -77.8966 | Allopatric |

| | | | | | | | |
|---|---|---|---|---|---|---|---|
| 77 | COR | Site5 | Site 5_PC_4-1 | RL-S | 34.37725 | -77.8966 | Allopatric |
| 91 | COR | Site5 | Site 5_PC_4-1 | RL-S | 34.37725 | -77.8966 | Allopatric |
| 20 | LAC | SpinKY | Spindletop_L_KY-c2 | BL-S | 38.01565 | -84.5052 | Allopatric |
| 70 | LAC | SpinKY | Spindletop_L_KY-c2 | BL-S | 38.01565 | -84.5052 | Allopatric |

4
